# Marine Bacteria Chemotaxis to Crude Oil Components with Opposing Effects

**DOI:** 10.1101/2022.01.26.477900

**Authors:** Xueying Zhao, Roseanne M. Ford

**Author notes:** **Correspondence** Roseanne M. Ford, Department of Chemical Engineering, University of Virginia, 102 Engineers’ Way, Charlottesville, VA 22904.

## Abstract

Marine microorganisms were critical to hydrocarbon removal from the Gulf of Mexico oil spill. Chemotaxis, a process attracting motile bacteria toward higher hydrocarbon concentrations, can increase biodegradation efficiency. However, crude oil also contains heavy metal ions that repel bacteria. Will bacteria migrate toward hydrocarbons or away from heavy metals? We exposed a marine isolate *Halomonas* sp. to decane and copper ions in a microfluidic device that maintained a constant concentration gradient across a channel. Bacterial distributions were used to quantify parameters in a mathematical model capturing bacteria motility and chemotaxis. This multi-scale model was adapted from the signal transduction mechanism of *E. coli*. For *Halomonas* sp., we used independent receptors for sensing attractant or repellent and chemotaxis parameter values were assessed. Predictions based on the multi-scale model correctly estimated the net attraction or repulsion responses of bacteria to the stimuli mixture. In some cases, the model yielded a stronger repulsion than what was observed experimentally, but still captured the general trends of bacteria distribution. Understanding how marine bacteria integrate information from multiple inputs to yield net migration toward or away from oil will improve predictions of hydrocarbon degradation rates.

## 1. Introduction

The largest oil spill in U.S. waters, the *BP Macondo* well blowout released around 780,000 m^3^ of crude oil into the Gulf of Mexico (Crone et al., 2010). To clean up the oil spill, processes such as burning, filtering, collecting, and dispersing were used. During the dispersion process, 1.8 million gallons of chemical dispersants were released into the Gulf of Mexico. Dispersants break the oil into small droplets which provides greater surface area for bacteria to be drawn to. However, the use of dispersants is controversial because of the unknown impact of these chemicals on marine ecosystems. An alternative approach to degrade oil droplets is to take advantage of the natural-occurring marine bacteria, which is called bioremediation.

Bioremediation is a strategy that uses microorganisms to remove pollutants from contaminated sites. These organisms can use pollutants as a food source and degrade them to less toxic or nontoxic substances (Hoff, 1993, Das et al. 2011). King et al. (2015) studied the biodegradation process following the oil spill. They found that the bacterial density in a cloud of dispersed oil (10^5^ cells/mL) was higher than outside of the cloud (10^3^ cells/mL), indicating that marine organisms played a role in oil spill cleanup. In addition, they found genes for chemotaxis and motility were enriched and expressed in organisms isolated from this cloud. Therefore, processes that increase bacteria accumulation in the cloud of dispersed oil are expected to aid cleanup as greater numbers of bacteria have the capacity to degrade more oil droplets. Chemotaxis is one of these processes; bacteria swim to a higher concentration of chemicals they can use as food and energy sources. The chemical compounds attracting chemotactic bacteria are called chemoattractants, while compounds bacteria swim away from are called chemorepellents. *Escherichia coli* show positive chemotaxis to amino acids and aromatic compounds, while sulfides and inorganic ions cause negative chemotaxis (Pandey and Jain, 2002). Studies (Lanfranconi et al., 2003, Meng et al., 2017) also showed that bacteria exhibit positive chemotactic responses towards n-hexadecane. Most importantly, the presence of chemotactic genes in *Oceanospirillales* cells suggests that marine bacteria are chemotactic to hydrocarbons in an oil plume (Mason et al., 2012). Interestingly, marine bacteria show different chemotactic responses to different hydrocarbons. Toluene, a major component of crude oil, has been reported as a negative chemoeffector for several marine *Pseudomonas* strains (Young and Mitchell, 1973). They also reported that inorganic copper ions act as chemorepellents to marine *Pseudomonas* strains. In contrast, de Sánchez and Schiffrin (1996) concluded that copper ions were strong chemoattractant to marine *Pseudomonas* strain H36-ATCC. Therefore, the effect of copper ions appears to vary for different marine bacteria.

Most published field studies focus on soil bacteria chemotaxis, while chemotaxis for marine bacteria is not as widely studied. Furthermore, there are few quantitative studies of marine bacteria chemotaxis. Seymour et al. (2008) studied three marine bacteria chemotactic responses in the presence of five different chemoattractants. They used the chemotaxis index to quantify the responses. In addition to chemotaxis, motility is also important for marine bacteria because of the heterogeneous environment of the ocean and transient release of nutrient pulses (Stocker and Seymour, 2012). The mean swimming speed for marine bacteria varies between 45 to 230 *μ*m/s based on observations of natural communities and isolates (Mitchell et al., 1995), while the swimming speed for *E. coli* is typically between 15 to 30 *μ*m/s. This higher swimming speed allows marine bacteria to quickly navigate around the temporal changes in nutrient concentration and enhance their likelihood of survival. Based on the diffusion profile of a population of bacteria, the random motility coefficient μ[m^2^/s] can be obtained in the absence of chemoeffectors. de Sánchez and Schiffrin (1996) reported the apparent random motility coefficient value of 1.5 × 10^−5^ cm^2^/s for marine *Pseudomonas* strain H36-ATCC. The lack of other quantitative studies prompts us to quantify chemotactic and motility parameters of marine bacteria.

In addition to quantifying marine bacteria chemotaxis to specific crude oil components, we also studied the chemotactic response to multiple stimuli because bacteria in natural settings are exposed to multiple compounds. When bacteria receive conflicting information from chemical signals, they decide how to respond and adjust their swimming behavior accordingly. At the individual bacterium level, chemotactic swimming is modulated by the directional change of rotation of the bacterium’s flagella. Figure S1 in Supporting Information shows the flagellar arrangement of *Halomonas titanicae* KHS3 in electron micrographs. *Halomonas titanicae* KHS3 is motile with peritrichously arranged flagella. The chemotactic mechanism of *Halomonas* sp. at the individual cell level is not well understood, while the molecular pathway of the chemotactic mechanism for *E. coli* bacteria has been studied extensively. For *E. coli*, when flagella rotate clockwise, bacteria tumble, while the counterclockwise rotation of flagella leads to a run along a straight pathway. In the absence of chemoattractant, bacteria trace out a random walk similar to diffusive motion (Rivero et al., 1989). When swimming through a higher concentration of attractant, chemotactic bacteria decrease their tumble probability (Tso and Adler, 1974), which results in a biased random walk that favors the attractant direction.

Several researchers (Mowbray and Koshland, 1987; Strauss et al., 1995; Kalinin et al. 2010; Zhang et al., 2019) have studied bacterial chemotactic responses in the presence of multiple stimuli. Middlebrooks et al. (2021) utilized a multi-scale model to predict *E. coli* responses to multiple stimuli *α*-methyl-DL-aspartate (MeAsp) and nickel ion. Kalinin et al. (2010) studied *E. coli* chemotaxis to multiple chemoattractants MeAsp and serine. They found when the ratio of receptors (Tar/Tsr) is greater than two, cells changed their preference from serine to MeAsp. For *Halomonas titanicae* KHS3, three methyl-accepting chemotaxis proteins are included in a chemosensory cluster (Gasperotti et al., 2018). It is possible for *Halomonas* sp. to use separate receptors to sense different compounds, e.g., receptor 1 for attractant and receptor 2 for repellent. Therefore, it is important to understand how the signal transduction mechanism inside an individual bacterium integrates an individual bacterium’s response in the presence of multiple stimuli and then relate it to the migration behavior of a bacterial population.

In order to measure the chemotactic response quantitatively, it is critical to create a well-defined chemical gradient since the chemical gradient is the driving force for chemotaxis. Middlebrooks *et al*. (2021) used a stopped flow diffusion chamber to analyze bacterial chemotactic velocities for MeAsp and nickel ion. The device they used was able to create a step change in the chemical concentration for the bacteria, which then relaxed over time in a predictable way due to diffusion. In contrast, the device in Wang et al. (2015) maintained a constant chemical concentration gradient; their design balanced the pressure at opposite sides of the microchannel to eliminate convective flow in the cross channel. Stocker et al. (2008) also used microchannels to study the chemotaxis of marine bacteria *Pseudoalteromonas haloplanktis*. They measured and modeled bacteria population responses to nutrient patches and plumes. They found that the chemotactic migration of marine bacteria *P. haloplanktis* was ten times faster than *Escherichia coli*. However, they didn’t investigate intrinsic cell transport parameters quantifying bacteria motility and chemotaxis.

In this study, *Halomonas* sp. were exposed to chemoeffectors in a microfluidic device that created a constant gradient. Decane was used as a chemoattractant. In addition to hydrocarbons, metallic ions (such as nickel, iron and copper) exist in all crude oil types in trace amounts. Thus, copper ions were investigated as a chemorepellent. Experimental data was used to quantify transport parameters in the muti-scale mathematical model capturing bacterial motility and chemotaxis that was adapted from Middlebrooks et al (2021). This work enabled us to quantify the effect of marine bacterial chemotactic processes that facilitate accessibility of oil droplets to bacteria, further increasing degradation of dissolved hydrocarbons.

## 2. Methods and Model

### 2.1 Preparation of Bacteria Cultures

Marine bacteria *Halomonas* sp. Bead 10BA (with genomic sequence similar to that of *Halomonas titanicae* strain KHS3 (Gasperotti et al., 2018) was obtained from Doug Bartlett (Scripps Institute of Oceanography). The *Halomonas titanicae* isolate is known to have a peritrichous arrangement of several flagella (Gasperotti et al., 2018) similar in that regard to *E. coli*. A TEM image of *Halomonas titanicae* flagella is shown in Figure S1 in the Supporting Information. In each experiment, 100 µL of *Halomonas* sp. Bead 10BA frozen stock was cultured in 50 mL marine broth (Fisher Scientific, NY) with 100 µL decane added. Bacteria were cultured in a 250 mL Erlenmeyer flask on a Thermo Scientific Incubated Shaker (MaxQ4000) with a rotation rate of 150 rpm at 30 ^°^C.

Bacteria were harvested at an optical density of 1.20 as previous studies (Doug Bartlett’s lab, personal communication) suggested the highest motility at late logarithmic phase (measured in spectrophotometer, Molecular Devices, Spectramax 384 Plus) at 590 nm. Staining stock solution was prepared by dissolving the contents of one vial CFDA SE (Invitrogen, CA) in 90 μL of DMSO (Invitrogen, CA) to reach a concentration of 10 mM. Then the stock solution was diluted in 5 mL of 5% random motility buffer (100% RMB included 9.13 g/L Na_2_HPO_4_ (Fisher Scientific), 4.87 g/L NaH_2_PO_4_·H_2_O (Amresco), 0.029 g/L EDTA (Sigma, MO)). Bacteria were incubated in the working solution for at least 60 mins at 30°C, filtered twice to remove the dye, and then resuspended into 5% RMB to an optical density around 1.40. Before taking microscopy images, the motility of the bacteria was examined under a Zeiss 100/1.25 oil lens with Nikon microscope (Digital Sight DS-5Mc).

### 2.2 Chemoeffectors Used in the Experiments

Decane was selected as the chemoattractant of interest because it is a major component in crude oil. To assay for chemotaxis, a decane solution was produced at the solubility limit in water by adding the amount of decane calculated for the solubility level. The solubility of n-decane in water is 0.009 mg/L at 20 °C (Verschueren, 2001), which corresponds to a concentration of 0.06 μM. Interestingly, in salt water, the solubility increases by an order of magnitude to 0.087 mg/L at 20 °C (Verschueren, 2001). Thus, making it feasible to test the response of *Halomonas* sp. to different concentrations of decane. The recipe for synthesized sea water was from Doug Bartlett’s lab (personal communication): Trizma base 5 g/L, KCl 0.75 g/L, NH_4_Cl 1 g/L, MgSO_4_.7H_2_O 3.91 g/L, MgCl_2_.6H_2_O 5.08 g/L, CaCl_2_ 1.5 g/L, and NaCl 23 g/L. Then HCl was used to adjust pH to 7.5. For repellent, copper sulfate was chosen because copper is known to be a repellent for marine bacteria (Young and Mitchell, 1973). Copper sulfate at a concentration of 0.05 mM was tested at first, but no obvious response was observed. Therefore, higher concentrations of 0.5 mM and 2 mM were selected. Note that Grey and Steck (2001) reported *E. coli* growth on agar plates of LB medium and of LB medium containing 4 mM CuSO_4_ and observed no difference. Thus, we concluded that concentrations up to 2 mM Cu ions were not toxic for bacteria. Taher (2016) reported the copper concentration in crude oil to be in the range of 0.0066 mM – 0.125 mM. The copper concentrations used in our experiments is higher than the range in the crude oil. We chose these higher concentrations because bacteria exhibited significant responses to copper ions at higher concentrations. Then we fitted the experimental results to the model to obtain parameters and these parameters can be used to predict bacteria responses at lower copper ion concentrations.

### 2.3 Microfluidic Design and Microscopy

Marine bacteria *Halomonas* sp. and chemoeffector were introduced into the uniquely designed microfluidic device (Wang et al., 2015) to obtain bacteria distribution in the presence of stimuli. The microscopic images shown in Figure 2 provides an example of a cross channel in the device where one end can maintain a constant concentration of bacteria source and the other end can maintain a constant concentration of chemoeffector source. This constant chemoeffector source was achieved by the reservoir from the top layer of the device. The design also balances the pressure at both ends, so transport occurs only by diffusion in the cross channel. More detailed information on the device design is provided in the Supporting Information. As Figure S6 shows, bacteria and the chemoeffector source introduced into two different arms of the Y-shaped channel did not mix with each other demonstrating that the microfluidic design worked properly. Microscopic images were taken using a Zeiss 780 confocal microscope with 10x objective lens in the Keck Center for Cellular Imaging at the University of Virginia. More information on the image analysis process can be found in Supporting Information.

### 2.4 Mathematical Model

#### Bacteria transport processes at the population level

Species mass conservation equations were used to model bacterial transport in the cross channel of the microfluidic device. Following the work of Wang et al. (2015), a one-dimensional equation was justified as no convective flow in the cross channel was observed for fluorescein. Therefore, diffusion controlled the mass transfer between the reservoirs of the cross channel; the device was designed to eliminate convection because it prevents bulk fluid flow from entering the cross channel.

Equation 1 shows the governing equation for bacteria concentration *b* at a steady state condition in the presence of a chemoeffector

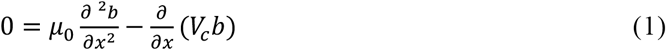

where *b* is bacteria concentration, *V*_*c*_ is the chemotactic velocity, the expressions of *V*_*c*_ are different under different cases and are detailed in the Supporting Information.

Equation 1 was solved using MATLAB R2018b to obtain bacterial distributions. Then least square regression analysis was used to fit Equation 1 to experimental data from steady state conditions in the presence of chemoeffectors to obtain chemotaxis parameter values. The plots were also generated in MATLAB R2018b.

#### Signal transduction kinetics

In order to understand how an individual bacterium reacts to chemoeffectors, we need to understand the bacteria signal transduction mechanism. The signaling response in *E. coli* chemotaxis depends on the phosphotransfer between a histidine kinase and a response regulator (Sourjik et al., 2010). Without the presence of any chemoeffector, the kinase CheA undergoes an autophosphorylation reaction, and then transfers the CheA phosphoryl group to the response regulator CheY. The signal is then transmitted to the flagellar motor, increasing the probability of clockwise (CW) rotation and results in bacteria tumbling. For *Halomonas titanicae* KHS3, Gasperotti *et al*. (2018) found that three methyl-accepting chemotaxis proteins are included in chemosensory cluster. However, the chemotactic mechanism for *Halomonas titanicae* is still unknown. Because *Halomonas titanicae* have the same flagella arrangement as that of *E. coli*, we first assumed they have the same chemotactic mechanism. However, the competitive model for the multiple stimuli did not fit the experimental results of *Halomonas* sp. We then assumed that chemotactic responses of *Halomonas* sp. to decane and copper act independently. More specifically, we assumed that each receptor regulates the response for either attractant or repellent as shown in Figure 1.

**Figure 1.**
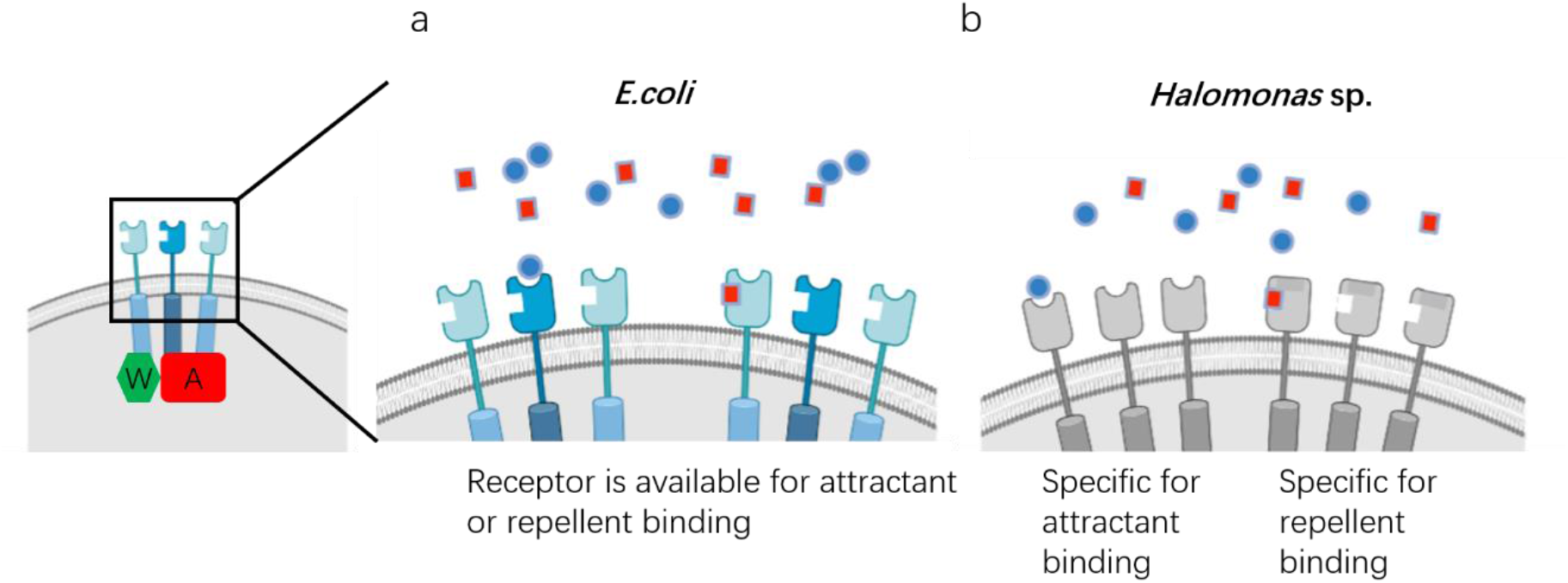
Schematic of the signal transduction mechanism in (a) *E. coli*, where receptor is available for both stimuli binding (blue indicates attractant and red indicates repellent); (b) *Halomonas* sp., where there are specific receptors for sensing attractant and repellent, separately.

Similar to the model for *E. coli*, in the absence of chemoeffector, we assume that there is an autophosphorylation reaction when the receptor complex is unbound. We assume that in the presence of a chemoattractant, binding of the attractant causes CheA inactivation, decreases the probability of clockwise (CW) rotation and causes the bacterium to tumble less frequently, which leads to the bacterium continuing to move in the same direction. In the opposite case, the binding of repellent does not inactivate CheA, and instead may enhance CheA activity, which increases the probability of clockwise (CW) rotation and results in more frequent tumbles, causing bacteria to swim away from the repellent source. We assume that decane binds specifically to one type of receptor and copper ion binds specifically to another type of receptor as shown in Figure 1b. This assumption is consistent with the response of bacteria to attractant and repellant being independent to each other, so the response to the chemoeffector mixture is a simple addition of the responses to attractant and repellent.

We further related the receptor concentration to the tumbling probability (more details are provided in the Supporting Information) and obtained Equation 2 for the chemotaxis velocity in the presence of decane:

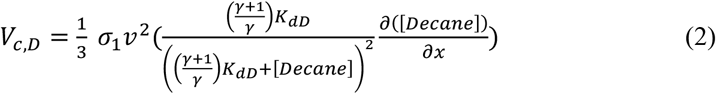

where *σ*_1_ is the stimuli sensitivity coefficient for attractant decane, *v* is the bacteria swimming speed, *γ* is the signaling efficiency, *K*_*dD*_ is the dissociation constant of decane, [*Decane*] is attractant decane concentration.

Note that Equation (2) simplifies to the same form as the chemotactic velocity equation in Wang et al. (2015) given that i) the ratio 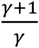 approaches 1 (*i*.*e. γ* ≫ 1) and ii) *χ*_0_ = *σ*_1_ *v*^2^, where *χ*_0_ is the chemotactic sensitivity coefficient, accounting for the strength of the chemotactic response for a population of bacteria. Different terms dominate the equation when the chemoeffector concentration is low or high relative to its binding dissociation constant. Given that *K*_*dD*_ was assumed to be at least one order of magnitude higher than the decane concentrations used in this study, the absolute concentration of decane did not affect the chemotactic velocity; the only effect of decane was through the increased magnitude of the decane concentration gradient.

## 3. Results and Discussion

*Halomonas* sp. were exposed to decane and migrated toward higher concentrations of it, which was consistent with decane being a chemoattractant. Upon exposure to copper, *Halomonas* sp. migrated away from higher concentrations of it, which was consistent with copper being a chemorepellent. Chemotactic parameters were obtained from attractant only and repellent only conditions under steady state. Then these parameters were incorporated into the multi-scale model to predict the migration direction of *Halomonas* sp. for competing scenarios with different chemical concentrations. Bacterial distribution profiles were compared with experimental results.

### *3*.*1 Halomonas* sp. chemotactic response to decane

A suspension of CFDA SE labeled *Halomonas* sp. was provided as a constant source at one end of the channel; decane, at the solubility limit of 0.06 µM, was provided at the opposite end. This scenario created a constant decane concentration gradient of 0.04 *μ*M/mm. Microscopic images of the cross channel were taken after the system reached steady state (at least two hours), as shown in Figure 2a. The distribution of chemotactic bacteria followed a curved parabolic shape with positive deviation from the diagonal line, which represents the control case when no chemoeffector was provided. This positive deviation indicates that decane is an attractant to *Halomonas* sp. A multi-scale mathematical model for transport of chemotactic bacteria was used to predict bacteria distribution by solving the governing equation (Equation 1) for bacteria and incorporating the chemotactic velocity for decane. (Equation 2). We collected experimental data for bacteria responses to two different concentrations of attractant (0.06 *μ*M and 0.2 *μ*M decane) to obtain parameter values *γ* and *σ*. As there were no literature values of *γ* and *σ* for *Halomonas* sp., values of these parameters for *E. coli* were used as the initial estimates. For the dissociation constant *K*_*dD*_, a value of order of magnitude 10^−6^M was used because we assumed this value was close to the maximum availability value (or the solubility level) in the water. In addition, a study by Xie *et al*. (2015) also reported a dissociation constant of order 10^−6^ M for marine bacteria chemotactic response. Parameters used in the model were thus the stimuli sensitivity coefficient for attractant σ_1_ = 6 s, signaling efficiency *γ* = 4, and the decane binding dissociation constant *K*_*dD*_ = *μ*M (as summarized in Table 1).

**Figure 2.**
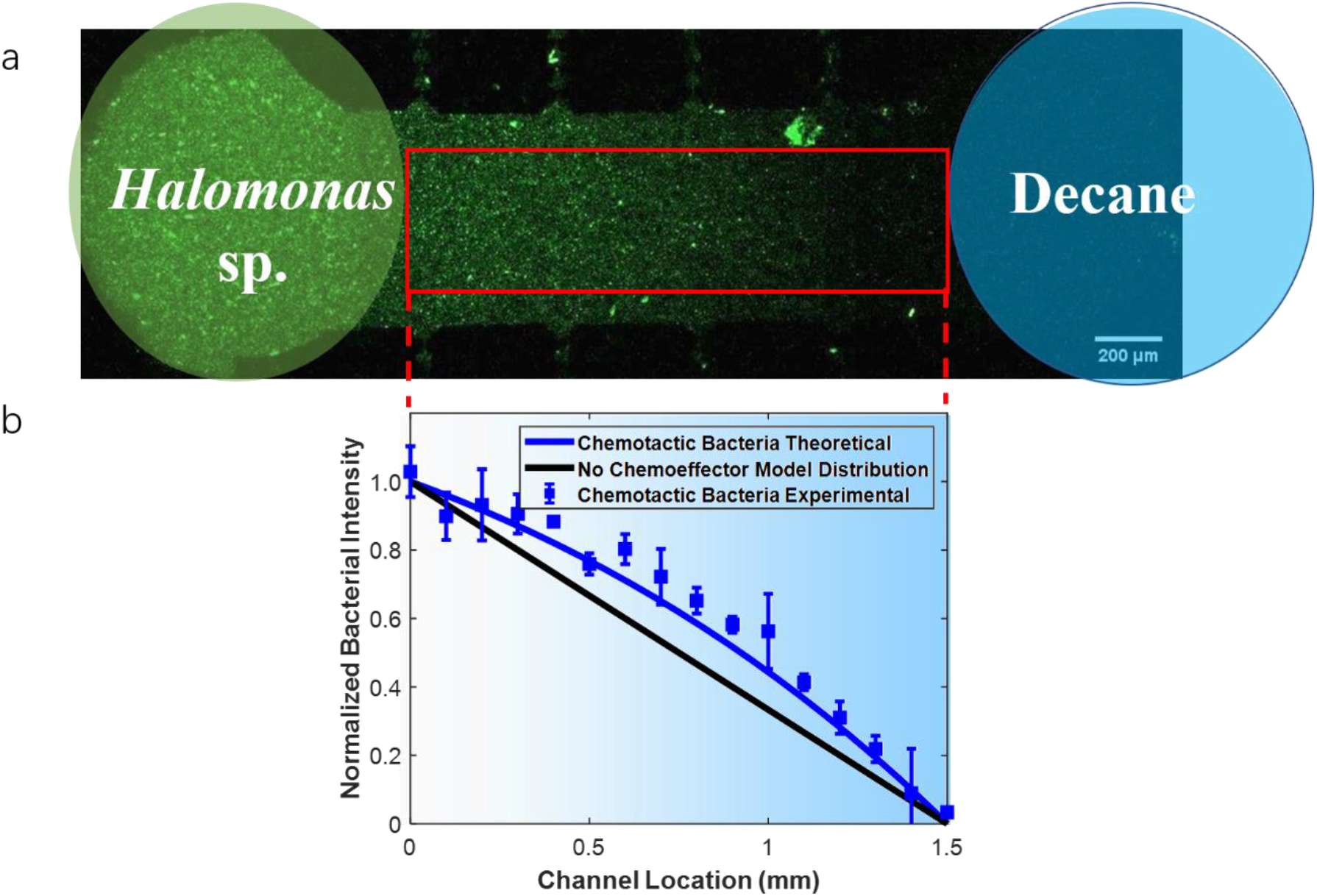
(a) Microscopic images of CFDA SE labeled *Halomonas* sp. distribution in a cross channel at steady state, where bacteria migrate from a constant source at the left-hand side toward 0.06 μM decane from a constant source at the right-hand side; the fluorescence intensity is proportional to bacterial concentration. The red rectangular box indicates the region over which the data was analyzed. (b) Normalized bacterial intensity profile from the experiment (blue squares) and model results (solid blue line). The black line indicates the expected bacterial distribution at steady state for the control case without chemoattractant. The blue shading indicates the concentration of decane which decreases from right to left. Errors bars represent standard deviation for two replicate experiments.

**Table 1.** Comparison of parameter values for Halomonas sp. and E. coli

| Symbol | Parameter | *Halomonas*<br>sp. | *E. coli* |
| --- | --- | --- | --- |
| $\gamma$ | Signaling efficiency | 4 | 4<sup>[1]</sup> |
| $\sigma_1$ | Stimuli sensitivity coefficient for<br>attractant (s) | 6 | 11<sup>[2]</sup> |
| $\sigma_2$ | Stimuli sensitivity coefficient for<br>repellent (s) | 3 | 11<sup>[2]</sup> |
| $\kappa$ | Repellent sensitivity coefficient | 2.7 | 9<sup>[1]</sup> |
| $v$ | Swimming speed ($\mu\text{m/s}$) | 50 | 22<sup>[3]</sup> |
| $\mu_0$ | Random Motility Coefficient<br>($\text{m}^2/\text{s}$) | $2.5 \times 10^{-10}$ | $1.5 \times 10^{-9}$<sup>[4]</sup> |

[1] Zhao et al., 2022; [2] Middlebrooks et al., 2021; [3] Wang, 2013; [4] de Sánchez and Schiffrin, 1996.

### 3.2 *Halomonas* sp. chemotactic response to copper ion

To assay bacterial chemotaxis to a chemorepellent, we followed a similar experimental setup that was used for the attractant decane. A suspension of CFDA SE labeled *Halomonas* sp. was provided at one end of the cross channel, and a solution of CuSO_4_ at the opposite end. This configuration created a constant gradient with a source of either 0.5 mM or 2 mM CuSO_4_. Microscopic images of the cross channel were recorded after the system had reached steady state. The images were then processed and analyzed to generate bacterial distribution data, as shown in Figures 3d and 3g. Distribution data for both concentrations follow a curved parabolic shape with negative deviation from the control case (when no chemoeffector was provided). These distributions indicated that Cu^2+^ was a repellent for *Halomonas* sp. as anticipated. Bacteria distribution profiles were also generated by solving the governing equation (Equation 1) for bacteria and using the chemotactic velocity derived for Cu^2+^ (Equation S13). We used the same value of *γ* from the result in the previous section for *Halomonas* sp. Values of κ = 2.7 and stimuli sensitivity coefficient for repellent σ_2_ = 3 s were obtained by fitting the model to data from the repellent experiments. As an initial estimate for regressing the parameter κ, we used a value of 9 that was previously determined for *E. coli* and nickel ion (Zhao et al., 2022). For the dissociation constant *K*_*dC*_, a value on the order of 10^−3^ M was used because this value corresponded to the concentration where bacteria exhibited the greatest repulsion from Cu^2+^. We also expect *K*_*dC*_ to be near concentrations associated with toxicity to bacteria. The value of *K*_*dC*_ = 3.5 mM was obtained after regressing the model predictions with the experimental results.

**Figure 3.**
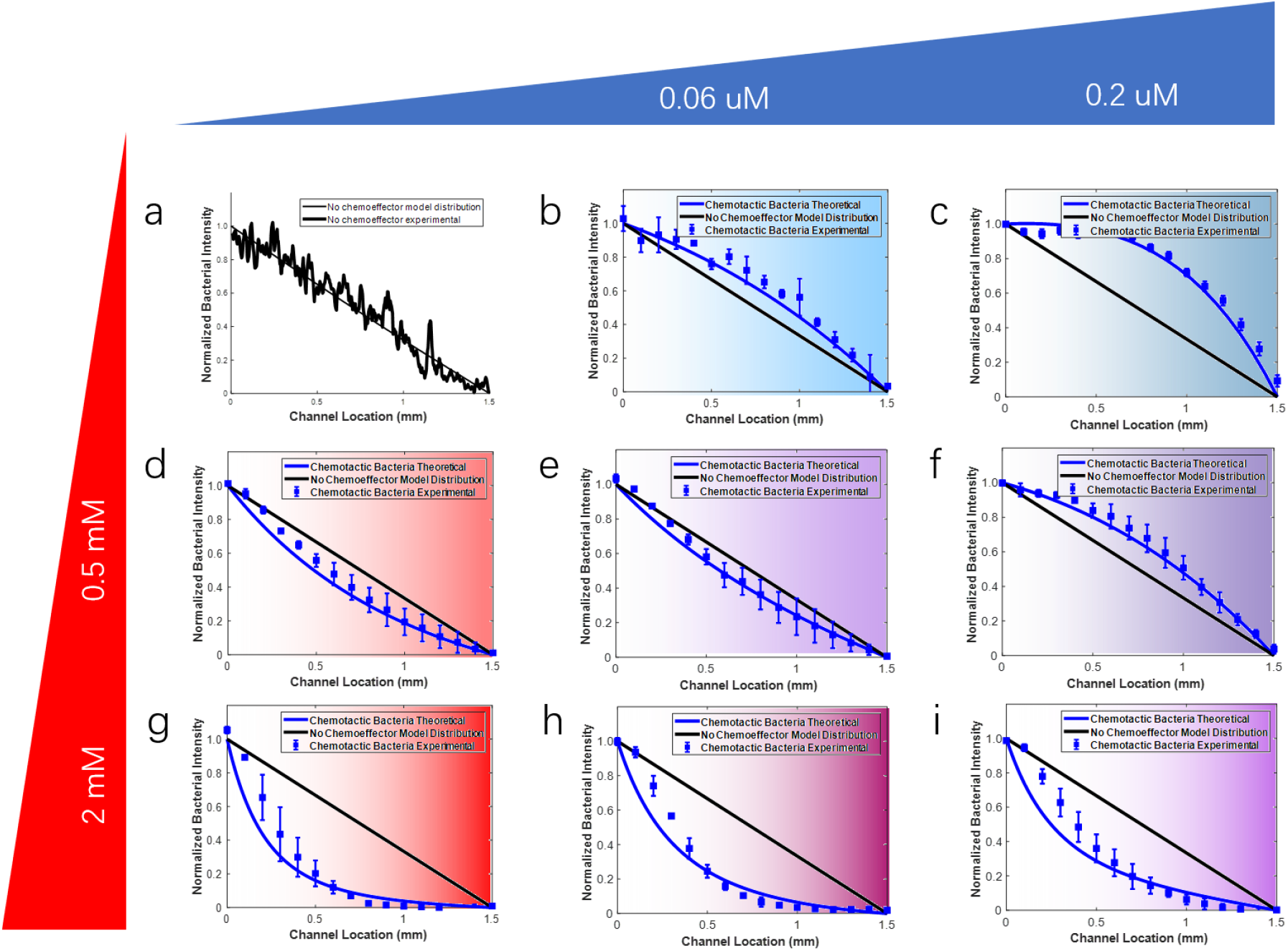
CFDA SE labeled *Halomonas* sp. distributions in a cross channel at steady state. There is a constant source of bacteria on the left-hand side of the channel and a constant source of buffer, attractant and/or repellent on the right-hand side: (a) buffer only as a control, (b) 0.06 μM attractant decane, (c) 0.2 μM attractant, (d-f) a combination of 0.5 mM repellent copper ions and 0, 0.06 or 0.2 μM attractant, respectively, (g-i) a combination of 2 mM repellent and 0, 0.06 or 0.2 μM attractant, respectively. The fluorescence intensity is proportional to bacterial concentration. The color shading indicates the concentration of chemoeffector which decreases from right to left. Errors bars represent standard deviation for (b-e) and (h-i) two replicate experiments, and (f) three replicate experiments.

### 3.3 *Halomonas* sp. response to chemoeffector mixtures

CFDA SE labeled *Halomonas* sp. were provided as a constant source at one end of the channel. Different concentration combinations of attractant decane (0.06 *μ*M and 0.2 *μ*M) and Cu^2+^ (0.5 mM and 2 mM) were provided at the opposite side. This arrangement provided constant gradients of chemoeffectors in the mixture. Figure 3 shows the normalized population density of *Halomonas* sp. in response to chemoeffector mixtures once the system reached steady state (after at least two hours). It shows that bacterial distributions for the various chemoeffector mixtures fall somewhere between that for attractant only and repellent only cases. This result suggests that bacteria integrate the signal from both chemoeffectors. By comparing Figure 3c and Figure 3f, we can see that the addition of Cu to 0.2 *μ*M decane decreased the migration of *Halomonas* sp. As the concentration of Cu increased from 0.5 mM to 2 mM, the bacteria response shifted from attraction to repulsion for the chemoeffector mixture, as shown in Figures 3f and Figure 3i. Bacteria distribution profiles were also predicted by solving the governing equation (Equation 1) for bacteria and using the chemotactic velocity of Equation S14 for the chemoeffector mixture. The values of *γ*, σ_1_, σ_2_, *κ* for *Halomonas* sp. from the results in the previous sections were used directly without additional fitting.

The effect of addition of decane to Cu was also shown in Figure 3. A comparison of Figure 3d and Figure 3e shows that the addition of decane to 0.5 mM Cu increased the migration of *Halomonas* sp. to the chemoeffector mixture. As the concentration of decane increased from 0.06 *μ*M to 0.2 *μ*M as shown in Figure 3f, *Halomonas* sp. switched the response from repulsion to attraction to the chemoeffector mixture. The model correctly predicted the response both quantitatively and qualitatively. In summary, the mathematical model was used to predict *Halomonas* sp. chemotaxis response to chemoeffector mixtures. Experimental results and model predictions for different scenarios are shown in Figure 3. The model predicted the experiment results successfully except for the cases with 2 mM Cu. For the highest Cu ion concentration (2 mM) the model predicted a greater deviation from the diagonal line than what was observed experimentally. The deviation from the diagonal line is a measure of the chemotactic response. Thus, our model appears to over predict the degree of repulsion of the bacterial population due to the Cu ion concentration. We suspect at the high Cu ion concentration the response may be limited by the number of available receptors or by another internal process in the chemosensory pathway that was not explicit in our multi-scale model. However, the copper concentrations in the crude oil are in the lower range of 0.5 mM, and our model clearly fit the data well in that range. Therefore, our model was consistent with experimental data over the copper concentration range of interest for oil spill bioremediation.

In conclusion, a mathematical model was presented in this paper to capture *Halomonas* sp. chemotactic response to attractant and repellent. The model was used initially to fit bacteria responses to single stimuli and obtain parameter values for *γ, σ*_1_, *σ*_2_, *κ, K*_*dD*_, *K*_*dC*_. Then the model was used to predict the response of *Halomonas* sp. to mixtures of decane and Cu^2+^. Note that in our model, we didn’t consider the adaptation of the chemotactic response, which is modulated by the methylation of the transmembrane receptors. Receptors are reset to baseline level through this methylation reaction if there is a significant increase in the local chemoeffector concentrations. However, the chemoeffector concentration is a continuous and gradual change in our experimental system since bacteria travel in a relatively shallow gradient. Therefore, there was no need to specify the methylation reaction in the model because CheA activation by receptor binding and CheR/CheB activities tended to balance out (Lele et al., 2015).

Overall, the model predictions captured *Halomonas* sp. responses well at lower Cu concentration near 0.5 mM. Specifically, for the case with 0.06 *μ*M decane and 0.5 mM Cu, *Halomonas* sp. our model predicted migration toward the mixutre, meaning chemotaxis will bring more bacteria to the mixture and likely increase biodegradation of hydrocarbons. We further applied the model to predict bacteria responses to chemoeffector concentrations found in crude oil (Taher, S. R., 2016), as shown in Figure 4. In all the scenarios, bacteria were attracted to the chemoeffector combinations. This outcome supports observations from the Gulf of Mexico oil spill that marine bacteria accumulated near the crude oil rather than being repelled away. Therefore, we assert that chemotaxis played an important role in biodegradation. The multi-scale model can be used to predict naturally-occurring marine bacteria responses to oil mixtures and provide insight on whether other interventions beyond bioremediation are needed to clean up oil spills.

**Figure 4.**
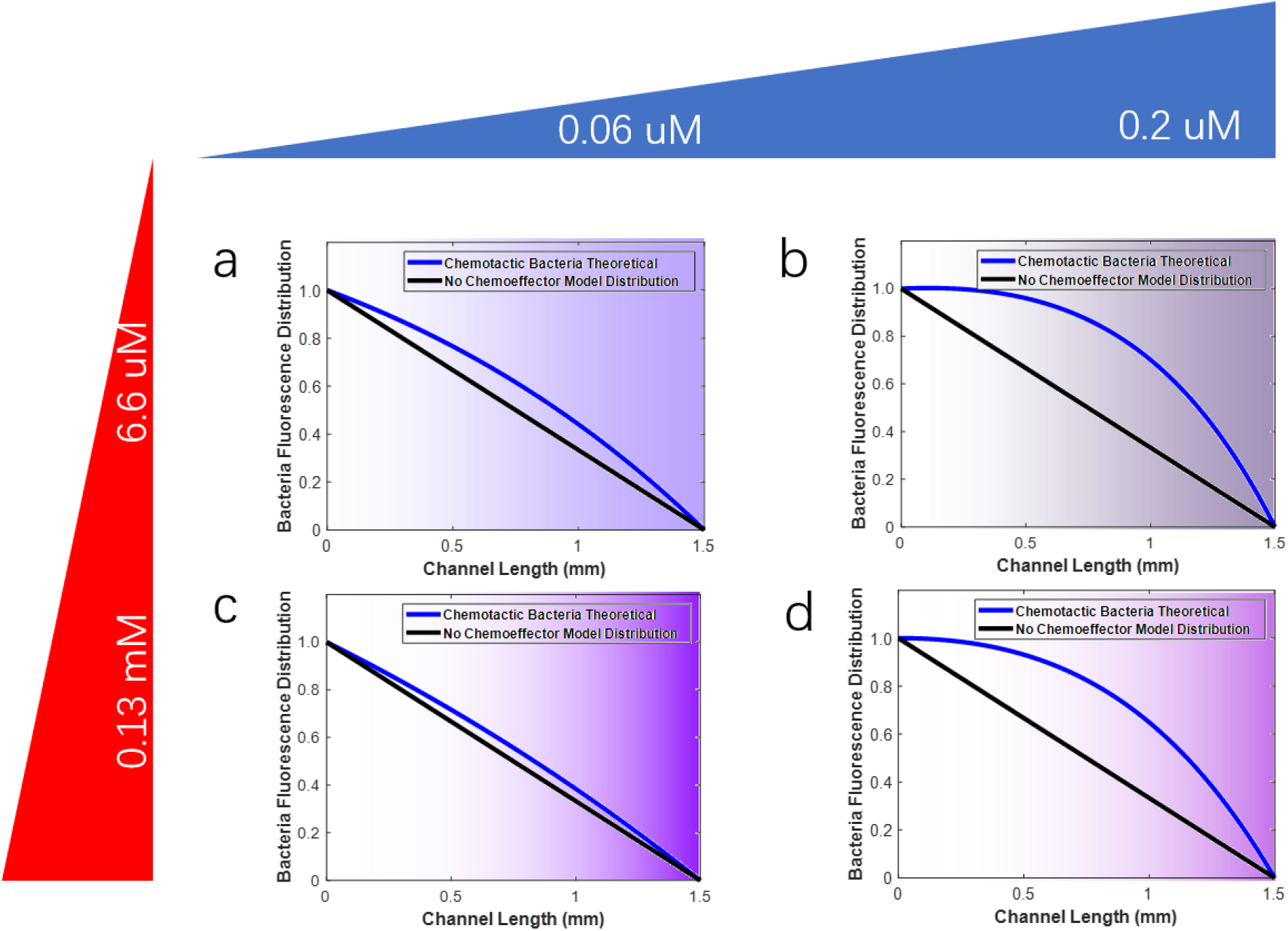
Prediction of *Halomonas* sp. distributions in a cross channel at steady state using the parameter values obtained in the study. (a) a combination of 0.06 μM attractant decane and 6.6 μM repellent copper ion, (b) a combination of 0.2 μM attractant and 6.6 μM repellent, (b) a combination of 0.06 μM attractant and 0.13 mM repellent, (b) a combination of 0.2 μM attractant and 0.13 mM repellent. The color shading indicates the concentration of chemoeffector which decreases from right to left.

## Acknowledgements

We appreciate Dr. Doug Bartlett from Scripps Institute of Oceanography for sharing *Halomonas* sp. Bead 10BA with us. We acknowledge the Keck Center for Cellular imaging for the usage of the Zeiss 780 confocal microscopy system to acquire the images (PI: AP; NIH OD016446). We thank Dr. Xiaopu Wang for discussing about microfluidic device, members in Dr. James P. Landers’ lab for helping with plasma treatment and members in Dr. T. Brent Gunnoe’s lab for helping with decane concentration measurement. This research was supported in part by a grant from Gulf of Mexico Research Initiative.

## Supporting Information

Additional supporting information may be found online in the Supporting Information section at the end of the article.

