## Supporting Information for "Marine Bacteria Chemotaxis to Crude Oil Components with Opposing Effects"

**Correspondence**

### Supporting Information

#### 1. SI Introduction

Figure S1 shows the flagellar arrangement of *Halomonas titanicae* KHS3. *Halomonas titanicae* KHS3 is motile with a peritrichous arrangement of flagella.

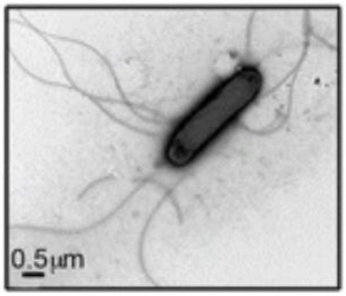


Figure S1. Transmission Electron Micrograph of *Halomonas titanicae* KHS3, showing flagellar arrangement. (From Gasperotti et al., 2018).

#### 2. SI Methods and Model

##### 2.1 Microfluidic design, fabrication and operation

The uniquely designed microfluidic device is made up of three layers (Wang et al., 2015). Polydimethylsiloxane (PDMS) was used to make the top and bottom layers. We chose PDMS because the oxygen permeability of PDMS allows the bacteria to maintain motility inside the channel. Both of the layers were treated with a plasma cleaner (PlasmaEtch, NV) to increase the hydrophilicity of PDMS. The top layer has two inlets connected to the main channel and the dimensions of the main channel are 85 μm high, 3.3 mm wide, and 2 cm long. The centerpiece was made from black polystyrene material

(Staples Inc.) (Zhao et al., 2022). A black centerpiece was used in order to block background fluorescence in the top channel from the signal in the cross channels in the bottom layer. Double-sided tape (3M) was used to adhere the centerpiece to the top and bottom layers of PDMS. We cut four pairs of vias into the centerpiece by laser cutter (VersaLaser, AZ) as depicted in Figure S2. The purpose of the vias is to connect the main channel in the top layer with the cross channels buried in the bottom layer. No convective flow will occur in the bottom channels because the pressures are equal in the vias at the cross positions perpendicular to the flow direction, thus diffusion is the only driving force in the bottom channel. The dimensions for the cross channels were 40 µm high, 600 µm wide, and 1.5 mm long.

To obtain the random motility coefficient of *Halomons* sp., 5% random motility buffer was provided at one end of the cross channel, while a bacterial suspension was provided at the other end of the channel. We replaced the buffer with chemoeffectors to study bacteria chemotaxis to an attractant and/or a repellent. The flow of bacteria and chemicals in the Y-shaped channel was controlled by a syringe pump SP 220i (World Precision Instruments, LLC, FL). The volumetric flow rate was set at 1.0 mL/h, corresponding to a linear speed of 2.42 mm/s. We calculated the Reynolds number (Re) in the main channel to be 0.17; for Re << 1 the two streams do not mix as they are under laminar flow.

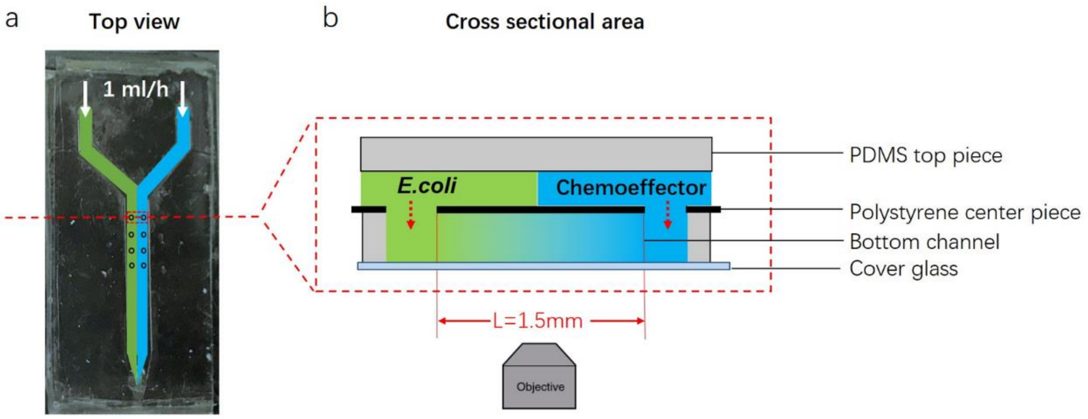


Figure S2. (a) Photo of microfluidic device (top-down view) and (b) an enlarged cross-sectional view. Adapted from Zhao et al., 2022.

#### 2.2 Microscopy and image analysis

A Zeiss 780 inverted confocal microscope with 10x objective lens was used to take microscopic images of bacteria distributions in the cross channel. The microscope was provided in the Keck Center for Cellular Imaging at the University of Virginia. A MBS 488/561 filter was used and the detector was in the range of 502-607 nm. The 1.5 mm-long cross channel was taken in two images, and then stitched together with overlap percentage 0.01% (Zhao et al., 2022). Five sequential images were collected for each region and superimposed together. For each image, we selected the region starting from the edge of the bacteria source via. We then measured the pixel number between scale markings and scaled the pixels to the known distance of 0.4 mm based on the channel design. A red rectangular box was drawn on the image to indicate the selected analysis region. Gray level values in each region were collected by using the plot profile feature.

The following equation was used to normalize the bacteria fluorescence intensity for each image:

$$N_j^n = \frac{N_j - N_0}{N_F - N_0} \tag{S1}$$

where $N$ represents the bacteria intensity, the superscript n represents the normalized value, subscript j corresponds to a certain location along the channel, $N_0$ is the bacteria intensity at the channel end opposite from bacteria input (*i.e.* sink), $N_F$ is the bacteria intensity at the channel end where fluorescent labeled bacteria are introduced (*i.e.* source).

#### 2.3 Mathematical Model

The governing equation for bacteria concentration *b* under unsteady state conditions without chemotaxis is

$$\frac{\partial \mathrm{b}}{\partial \mathrm{t}} = \mu_0 \frac{\partial^{\ 2} b}{\partial x^2} \tag{S2}$$

where $\mu_0$ is the bacterial random motility coefficient. The boundary conditions are $b\ (0, t) = b_0$ and $b\ (L, t) = 0$ and the initial condition is $b\ (x, 0) = 0$. The solution to Equation S2 for a channel of finite length *L* is

$$\mathrm{b}\ (x, t) = 1 - \frac{x}{L} - \frac{2}{\pi} \sum_{n=1}^{\infty} \frac{\sin n\pi \frac{x}{L}}{n} \exp(-\frac{n^2 \pi^2 \mu_0 t}{(L)^2}) \tag{S3}$$

We obtained the random motility coefficient $\mu_0$ by using nonlinear least square regression analysis to fit Equation S3 to experimental data from unsteady state conditions without the addition of chemoeffector.

We assumed that attractant and repellent bind independently to different receptors A and B. This assumption is the primary difference between the model for *Halomonas* sp. and *E. coli*. Phosphorylation reactions in the absence of chemoeffector were represented by the following equations:

$$A + P_i \overset{K_1}{\Leftrightarrow} A\text{-}P_i \quad (S4)$$

$$A_T = A + A\text{-}P_i \quad (S5)$$

$$B + P_i \overset{K_1}{\Leftrightarrow} B\text{-}P_i \quad (S6)$$

$$B_T = B + B\text{-}P_i \quad (S7)$$

$$R_T = A_T + B_T \quad (S8)$$

where $R_T$, a fixed value, is the summation of $A_T$ and $B_T$, $A_T$ is the summation of A, the unbound receptor complex for attractant binding, $B_T$ is the summation of B, the unbound receptor complex for repellent binding, A-$P_i$ is the phosphorylated receptor, and $K_1$ is the dissociation constant of phosphorylation of the unbound receptor complex. In the absence of any other evidence, we assigned the same dissociation constant value for the phosphorylation reactions of A and B.

In the presence of attractant and repellent, the receptor undergoes the following reactions:

$$A + D \overset{K_{dD}}{\Longleftrightarrow} D\text{-}A \quad (S9)$$

$$B + Cu \overset{K_{dC}}{\Longleftrightarrow} Cu\text{-}B \quad (S10)$$

$$Cu\text{-}B + P_i \overset{K_2}{\Leftrightarrow} Cu\text{-}B\text{-}P_i \quad (S11)$$

$$[R_T] = [A] + [B] + [D\text{-}A] + [Cu\text{-}B] + [A\text{-}P_i] + [B\text{-}P_i] + [Cu\text{-}B\text{-}P_i] \quad (S12)$$

In the above Equations S9-S12, D-A and Cu-B represent bound receptor complexes of decane and copper ions, respectively, $K_{dD}$ is the dissociation constant of decane, $K_{dC}$ is the dissociation constant of copper ions, and $K_2$ the dissociation constant of phosphorylation of the Cu ion bound receptor complex, and $R_T$ is the total amount of receptor and is a fixed number,

Following the steps in the Supplementary Information in Zhao et al. (2022), the chemotactic velocities in the presence of copper ion only and in the presence both decane and copper are shown in Equation S13 and Equation S14, respectively:

$$V_{c,C} = -\frac{1}{3}\sigma_2(\kappa - 1)v^2 \frac{(\frac{\gamma+1}{\gamma})K_{dC}}{((\frac{\gamma+1}{\gamma})K_{dC}+(\frac{\kappa+\gamma}{\gamma})[Cu])^2}\frac{\partial([Cu])}{\partial x} \quad \text{(S13)}$$

$$V_{c,T} = -\frac{1}{3}\ v^2[\sigma_2 \frac{\left(\frac{\gamma+1}{\gamma}\right)(\kappa-1)K_{dC}}{(\left(\frac{\gamma+1}{\gamma}\right)K_{dC}+\left(\frac{\kappa+\gamma}{\gamma}\right)[Cu])^2}\frac{\partial[Cu]}{\partial x} - \sigma_1 \frac{\left(\frac{\gamma+1}{\gamma}\right)KdD}{\left(\left(\frac{\gamma+1}{\gamma}\right)KdD+[Decane]\right)^2}\frac{\partial[Decane]}{\partial x}] \quad \text{(S14)}$$

where $\sigma_1$ is the stimuli sensitivity coefficient for attractant decane, $\sigma_2$ is the stimuli sensitivity coefficient for repellent copper ion, $v$ is bacteria swimming speed, $\gamma = \frac{K_1}{[P_i]}$ is the signaling efficiency, $\kappa = \frac{K_1}{K_2}$ is the repellent sensitivity coefficient, $K_{dC}$ is the dissociation constant of copper, $K_{dD}$ is the dissociation constant of decane, $[Cu]$ is the repellent concentration, and $[Decane]$ is attractant concentration.

It is obvious that Equation S14 is simply the addition of Equation 2 and Equation S13, which is consistent with our assumption that chemoeffector binding to chemotaxis receptor complexes is non-competitive.

#### 3. SI Results and Discussion

##### 3.1 *Halomons* sp. random motility coefficient

A constant source of CFDA SE-labeled chemotactic *Halomonas* sp. was maintained at one end of the cross channel and buffer was maintained at one other end of the cross channel in experiments evaluating the random motility coefficient (*i.e.* bacterial diffusivity). Figure S3a shows bacteria distributions at several times as bacteria migrated through the channel; Figure S3b shows the corresponding normalized bacteria intensity. Plots in Figure S3b followed the expected exponential decay in bacteria concentration from the source at x=0. At long times, once steady-state was achieved (~ 60 min), the normalized bacterial distribution reached a linear trend. A random motility coefficient of $\mu_0 = 2.5 \pm 0.22 \times 10^{-10}\ m^2/s$ was obtained by using nonlinear regression for a best-fit of the model (Equation S3, n=10) to experimental data (averaged over four replicates). Model curves were also plotted using $\mu_0 = 2.5 \times 10^{-10}\ m^2/s$ in Figure S3b for comparison with the experimental results.

The Boltzmann transformation was used to convert the spatial and temporal dependence into a single parameter for unsteady diffusive processes, as shown in Equation S15:

$$\frac{b}{b_0} = \mathrm{erfc}\left(\frac{1}{2\sqrt{\mu_0}}\varepsilon\right) \quad \text{(S15)}$$

where $\varepsilon = \frac{x}{\sqrt{t}}$ with units of $mm/\sqrt{s}$. This transformation allowed us to compare the experimental results and model predictions at different times. As shown in Figure S3b Inset, the normalized bacteria intensity with respect to $\varepsilon$ at different times overlapped onto a single curve, indicating a diffusive process as a population of bacteria migrates in the channel with $\mu_0 = 2.7 \times 10^{-10}\ m^2/s$.

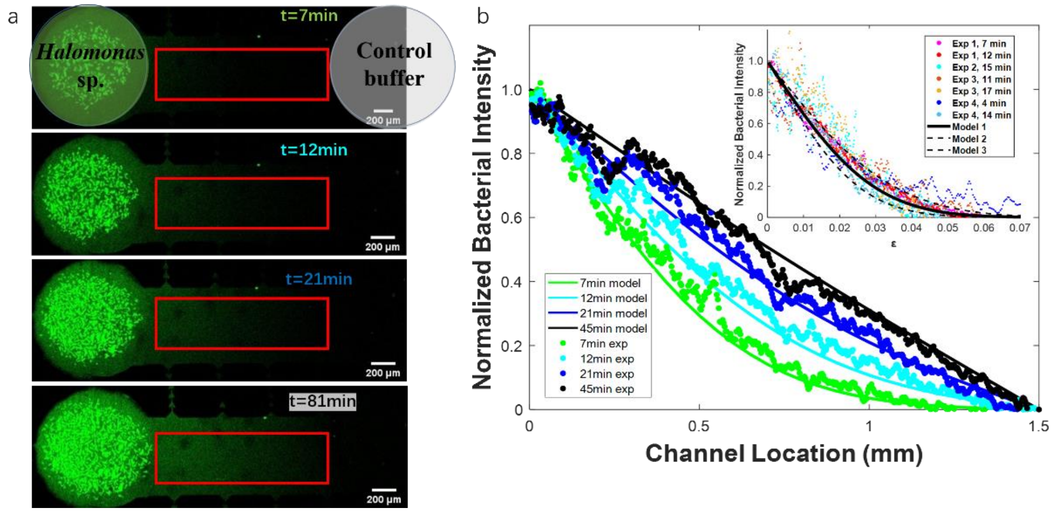


Figure S3. (a) CFDA-SE labeled *Halomonas* sp. distribution in a cross channel at several times. A constant source of bacteria was maintained on the left-hand side of the channel and a constant sink on the right-hand side. Bacteria concentration is proportional to the fluorescence intensity. The red rectangular box on each image indicates the area of interest where data was analyzed. (b) The comparison of normalized bacterial intensity along the channel (scattered data) and model predictions (solid lines) at several times. This set of data led to a random motility coefficient of $\mu_0 = 2.7 \times 10^{-10}\ m^2/s$. (*Inset*) Normalized bacteria intensity for four sets of experimental data collected at several times with respect to the Boltzmann transformation parameter $\varepsilon = \frac{x}{\sqrt{t}}$. Model 1 corresponds to the solid curve with $\mu_0 = 2.5 \times 10^{-10}\ m^2/s$, Model 2 and Model 3 corresponds to the upper dashed curve with $\mu_0 = 3.7 \times 10^{-10}\ m^2/s$ and the lower dashed curve with $\mu_0 = 1.9 \times 10^{-10}\ m^2/s$, respectively.

##### 3.2 Modulating *Halmonas* sp. response to a chemoeffector with increasing concentrations of an opposing stimulus

In this section we present microscopic images, normalized bacterial intensity distributions and model predictions for scenarios with different chemoeffector concentrations. Increasing Cu ion concentrations counter bacteria attraction to decane and change their response from overall attraction to repulsion, as shown in Figure S4. Similarly, increasing decane concentrations counter the repellent effect of Cu ion; altering the chemotactic response to the chemoeffector combination from overall repulsion to attraction, as shown in Figure S5.

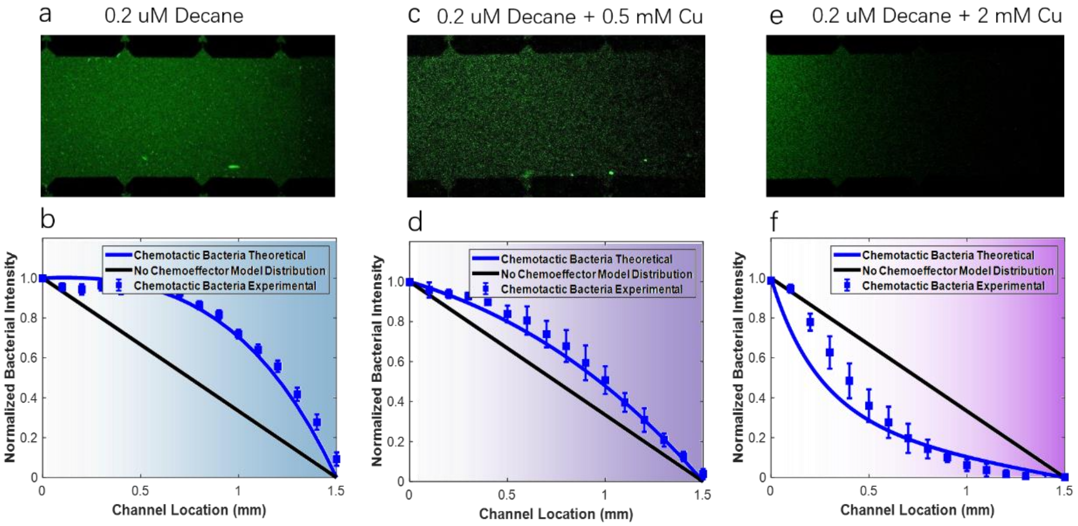


Figure S4. CFDA SE-labeled *Halomonas* sp. distribution in a cross channel at the steady-state. A constant source of bacteria was maintained on the left-hand side of the channel, and a constant source of chemoeffector was maintained on the right-hand side with concentrations: (a) 0.2 $\mu$M decane, (c) 0.2 $\mu$M decane and 0.5 mM Cu ions, and (e) 0.2 $\mu$M decane and 2 mM Cu ions. Bacterial concentration is proportional to the fluorescence intensity from the image and is plotted as normalized intensity as a function of location for bacteria response to (b) 0.2 $\mu$M decane, (d) 0.2 $\mu$M decane and 0.5 mM Cu ions, and (f) 0.2 $\mu$M decane and 2 mM Cu ions. Note that the gray level value in each image was calibrated itself using the approach described in the SI Methods and Model section, which eliminated fluctuation in the absolute value of fluorescence intensity for different images.

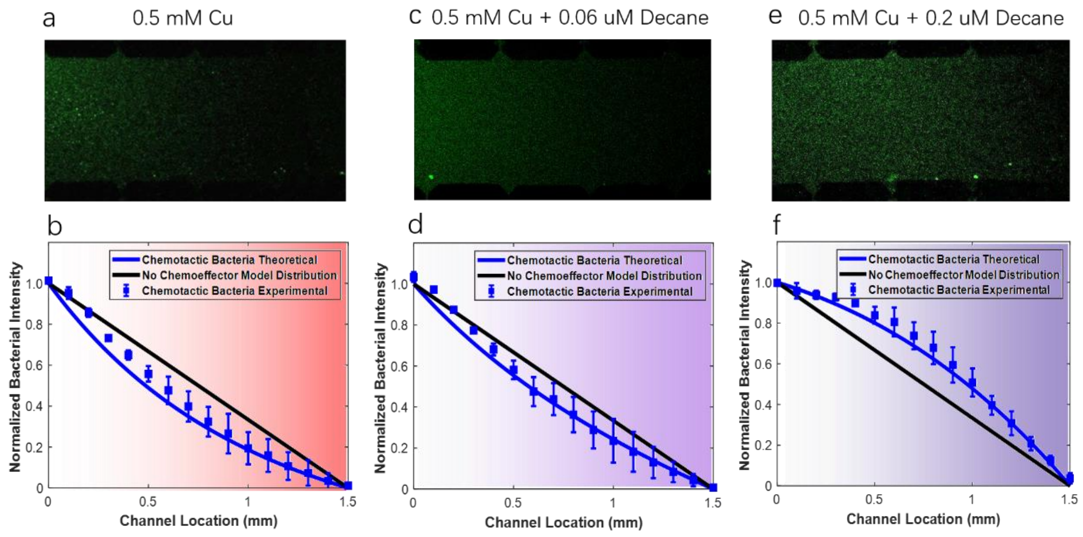


Figure S5. CCFDA SE-labeled *Halomonas* sp. distribution in a cross channel at the steady-state. A constant source of bacteria was maintained on the left-hand side of the channel, and a constant source of chemoeffector was maintained on the right-hand side with concentrations: (a) 0.5 mM Cu ions (c) 0.5 mM Cu ions and 0.06 $\mu$M decane and (e) 0.5 mM Cu ions and 0.2 $\mu$M decane. Bacterial concentration is proportional to the fluorescence intensity from the image and is plotted as normalized intensity as a function of location for bacteria response to (b) 0.5 mM Cu ions, (d) 0.5 mM Cu ions and 0.06 $\mu$M decane, and (f) 0.5 mM Cu ions and 0.2 $\mu$M decane. Note that the gray level value in each image was calibrated itself using the approach described in the SI Methods and Model section, which eliminated fluctuation in the absolute value of fluorescence intensity for different images.

##### 3.3 Bacteria adsorption on the PDMS top piece in the microfluidic device

In the process of disassembling the microfluidic device after our experiments we observed an interesting phenomenon. A noticeable band of high bacterial density was adsorbed to the PDMS top layer as depicted in Figure S6a. The intensity of the band increased with increasing Cu ion concentration. The zoomed-in images of the channel (as shown in Figure S6b) of the PDMS top piece demonstrated that the bacteria bands were brighter as the Cu concentration increased from 0.5 mM, to 2 mM and to 5 mM. Mixtures with decane appeared to reduce the band intensity in proportion to the decane concentration; adsorbed bands were not observed when decane was the sole chemoeffector. Figure S6c shows gray level values plotted across the channel from left to right using the Plot Profile feature in ImageJ. The peak location shifts away from the side where Cu was introduced. This peak shift is because bacteria tend to seek an optimum stimuli concentration, thus swimming further away from the Cu ion source as its concentration increases. The formation of a traveling wave (or band of high bacterial density) is consistent with predictions from mathematical models of bacterial chemotaxis. For example, a similar high density band was reported in Lanning et al. (2008) for bacteria suspended in the aqueous phase. What was different about our observation was bacteria adsorbed to the PDMS surface and retained a visual record of the distribution in the Y-shaped channel. We suspect the Cu ions increased the ionic strength, which enhanced adsorption of bacteria on the PDMS surface.

The adsorbed bands provided a nice confirmation that the microfluidic device was operating properly under laminar flow in the Y-shaped channel with very little mixing across the two streams. High density bands form in the fluid phase as bacteria sense and respond to chemoeffectors in the adjacent fluid stream and over the course of the experiment bacteria adsorb to the PDMS, reflecting the concentration in the fluid phase. The bands occurred along the center of the channel far from the vias that connected to the cross channels in the layer below ensuring that concentrations in the vias represented contents introduced in the appropriate stream.

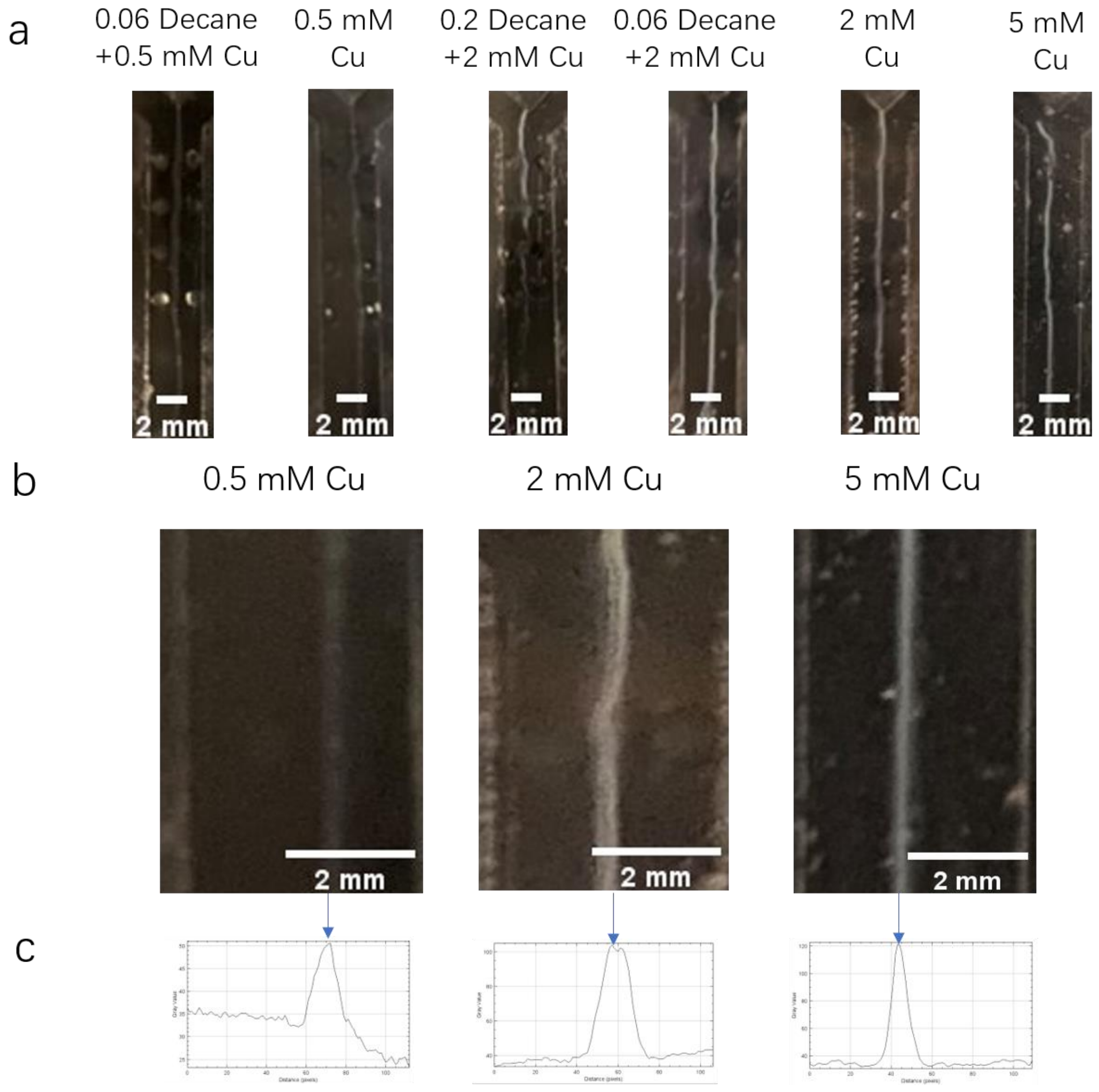


Figure S6. (a) Photos showing bacterial sorption above the Y-shaped channel for a section of the PDMS top piece after experiments were completed (around 2 hours) for different chemoeffector conditions. (b) Zoomed-in images of the PDMS top piece corresponding to experimental conditions of Cu ion concertation at 0.5 mM, 2 mM, and 5 mM. (c) Image gray level values plotted versus location across the channel for the corresponding images in (b). Blue arrows indicate the peak location.
